# Impact of body image on the kinematics of gait initiation

**DOI:** 10.1101/2023.11.11.566692

**Authors:** Kyosuke Oku, Shinsuke Tanaka, Yukiko Nishizaki, Chie Fukada, Noriyuki Kida

## Abstract

In our daily lives, we walk naturally because we can consider our physical characteristics and formulate appropriate motor plans. However, the impact of changes in body image during motor planning on walking movements remains poorly understood. Therefore, in this study, we investigated how walking movements change when body image is altered. We included 26 participants (13 males, 13 females, aged 18.27±0.52) to perform walking movements under five conditions: open eyes (baseline), closed eyes, closed eyes while imagining their bodies becoming large, closed eyes again, and open eyes again. As a result, under the condition where participants imagined their bodies becoming large, their stride length, step completion time, and foot lift height increased. These results are attributed to the disparity between actual body size and body image, which affects motor planning. The results of this study have potential applications in rehabilitation and sports coaching settings.

## Introduction

In our daily lives, we perform walking movements without conscious effort, especially when considering our own physical characteristics and the intended movements. Research suggests that this movement planning and movement imagery have equivalent functions [1-3]. Studies have indicated that when mentally imagining movement, the same brain regions are activated as when actually performing the movement [4]. Movement imagery is defined as “the mental expression of movement without actual bodily movement” [5]. Taking into account one’s physical characteristics is considered important in movement imagery.

Walking movements are believed to be planned by reverse calculation based on the desired speed. These movements are goal-oriented, involving lower-limb movements. During walking, step length and step frequency are adjusted to align with the walking goal at a specific speed [6]. Particularly, research has focused on the transition from a stationary state to the first step of walking [7-13]. Studies on walking initiation have shown correlations between electromyography readings of ankle muscles that control the leg lifting posture from the ground and forward walking speed [14]. Furthermore, studies on the center of pressure (CoP) of the body have indicated that the transition of CoP before walking initiation is related to forward walking speed [15-17]. Based on these previous studies, walking is planned based on the walking speed and executed while simultaneously controlling the center of gravity and the related muscles.

Motor imagery can encompass various types of movements, and walking movements are particularly useful during rehabilitation. Motor imagery has been noted as a useful approach for promoting motor function recovery after cerebrovascular disorders, particularly in relation to walking movements [18]. Motor imagery training, involving dorsiflexion and internal/external rotation of the hand joints, has been reported to improve voluntary control of paralyzed limbs in stroke patients [19]. In a study targeting individuals aged 65 years and above, the incorporation of motor imagery into physical training significantly improved balance ability, walking function, and self-efficacy in preventing falls [20, 21]. Thus, motor imagery is considered effective in improving walking movements.

Regarding the imagery of movement, it is necessary to consider one’s own physical characteristics. The recognition of one’s physical characteristics has been investigated using the concept of body image. Body image is defined as the conscious brain representation of the body, including self-image and appearance perception, which is derived from one’s posture perception [22]. The discrepancy between this body image and the actual physical condition is believed to affect motor performance. For example, in the case of walking, a decrease in stride length has been observed when stilts are used to extend the length of the lower limbs [23]. It is speculated that this is because the body image remains the same as the original body; however, the actual body has been extended, resulting in a failure to generate the necessary torque. Thus, the discrepancy between body image and actual physical condition may affect motor performance.

Visual information is believed to play an important role in manipulating body image. A person’s body image is thought to be changed by the height of the visible perspective and the presence or absence of visual information in the surrounding environment. Blocking visual information also has a significant impact on movement. Research has shown that walking with closed eyes results in longer stride length and longer time per step [24, 25]. Additionally, studies have indicated that raising viewpoints through virtual reality (VR) can make individuals perceive themselves as large, causing increase in their stride length [26]. Similarly, instructing individuals that their body has become large while raising the viewpoint through VR has also been shown to increase their stride length [27]. Thus, visual information plays a significant role in walking movements and posture control, and body image can have a significant impact on movement.

This study aimed to investigate the relationship between motor imagery and motor output by comparing the initial kinematics of walking in open-eyed, closed-eyed, and closed-eyed states with manipulated body images. Previous studies have shown that when the viewpoint is elevated using VR, individuals perceive their bodies as large, leading to an increase in their stride length [26]. In this study, we examined the effects of imagining a large body image using simple verbal instructions only during the initial phase of walking. Additionally, we aimed to clarify the mechanism of motor output by examining the kinematics of movements when body image is altered. We hypothesized that if the body image was larger than the actual body size, the stride length and height of lifting the feet would increase. By elucidating the motor output mechanism, the results of this study are anticipated to have applications in rehabilitation and sports

## Methods

### Participants

The experiment involved 26 healthy adults (13 males, 13 females, aged 18.27±0.52 years). Participants were allowed to use either their dominant or non-dominant feet. The sample size was determined based on previous studies (n=26) [28-32]. Prior to the experiment, the purpose and procedure were explained to the participants, and written consent for their participation was obtained. This study was approved by the Ethics Committee of Kyoto Institute of Technology and was conducted in accordance with the Declaration of Helsinki.

### Experimental setup and procedure

A starting point was marked with tape, and the participants were instructed to perform a 4-step walking motion barefoot. Two infrared cameras (OptiTrack and NeuralPoint) were placed approximately 6 m in front of the participants to capture their walking motions. Reflective markers, each with a diameter of 16 mm, were attached to the tips of the second toes of both feet to obtain three-dimensional coordinates. These infrared markers were captured using two infrared cameras, and data on the three-dimensional coordinates of the toes were obtained at a time resolution of 120 Hz.

The starting position was a standing posture with both toes aligned, and the participants were instructed to start walking from this position. They were asked to take four steps and then stop with both feet together. The foot used to initiate walking was not specified, and the participants’ dominant foot was not considered. First, the participants performed the walking task ten times under open-eye conditions. Subsequently, under closed-eye conditions, the participants were asked to wear an eye mask and perform the walking task ten times. After completing the task, they were asked to open their eyes and return to their starting point independently. Then, while wearing the eye mask, they were instructed to imagine themselves as if they had become large and to start walking. They were told to imagine that their body proportions remained the same but their size had increased to the point where their head would touch the ceiling of the laboratory (approximately 4 m). They performed the walking task ten times under this condition. Subsequently, as a post-test, they repeated the closed-eye condition three times and the open-eye condition three times. After the experiment, they answered a questionnaire to evaluate how well they were able to imagine themselves as if they had become large, using a 5-point scale.

## Data analysis

After analyzing the data of 20 participants who were able to imagine their body image well in the post-experiment survey, we focused only on the movement of the first step of walking. Time-series data of the 3D positions of the infrared markers attached to the tips of the second toes of both feet were preprocessed using a second-order Butterworth low-pass filter with a cutoff frequency of 10 Hz [33]. The start of the movement was defined as the onset when the speed continuously exceeded 30 cm/s for the first 5 frames; the end of the movement was defined as the offset when the speed continuously remained below 30 cm/s for 5 frames. This threshold was set at 10% of the maximum speed for all participants. The toe height and stride length, time taken for one step, were calculated for each trial, and the average values were determined for each condition.

## Results

The motion trajectory in the sagittal plane (Y-Z) is shown in Fig 1. The results indicated that when participants assumed a large body image, the stride length increased, and there was a possibility of higher foot elevation. Additionally, upon examining the motion trajectory, it became evident that, instead of following a parabolic curve, there is a possibility of performing a movement in which the foot is brought up to the landing point and then lowered straight down.

**Fig 1.**
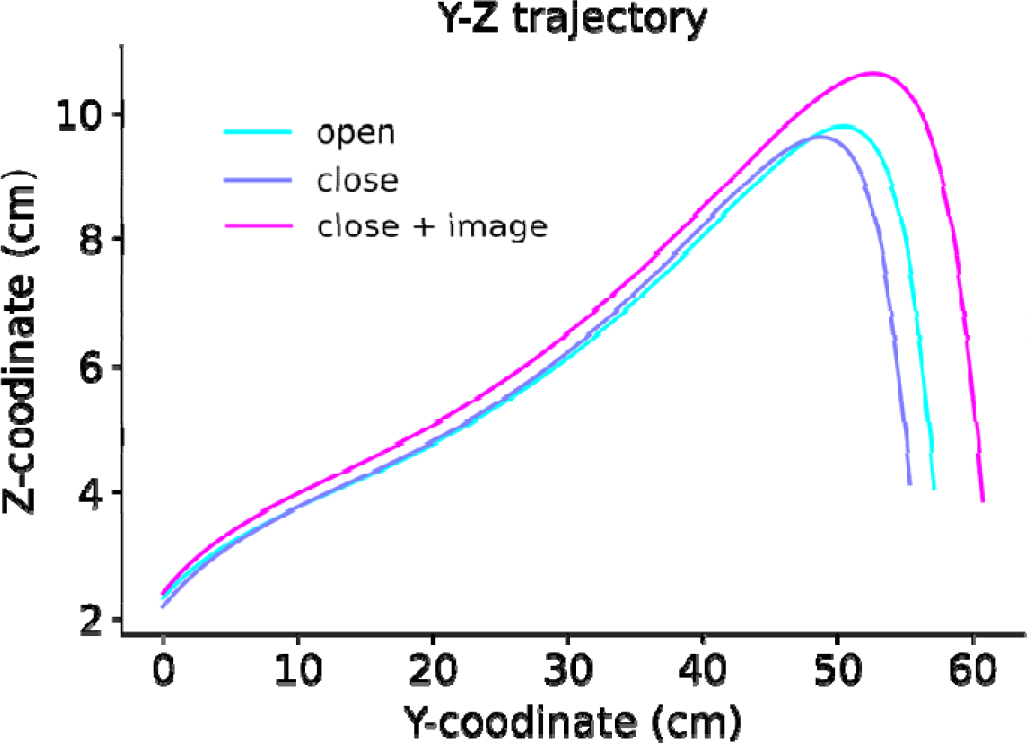
The trajectory of motion in the sagittal plane (Y-Z direction). The color of the lines represents different conditions, including the open condition, close condition, and close condition with imagined body enlargement (close + image) being depicted.

Fig 2 shows the average stride length across participants standardized by the lower limb length for each condition. A one-way analysis of variance was conducted with the conditions as the factors. The analysis revealed a significant main effect (*F* [4, 76] = 13.68, *p* = 1.90×10^-8^, η^*2*^ = 0.07). Subsequently, Bonferroni’s multiple comparisons test revealed significant differences between the closed eyes and the closed eyes + image condition, between the closed eyes + image and the second closed eyes condition, and between the post-closed eyes and the open eyes condition (p < 0.05).

**Fig 2.**
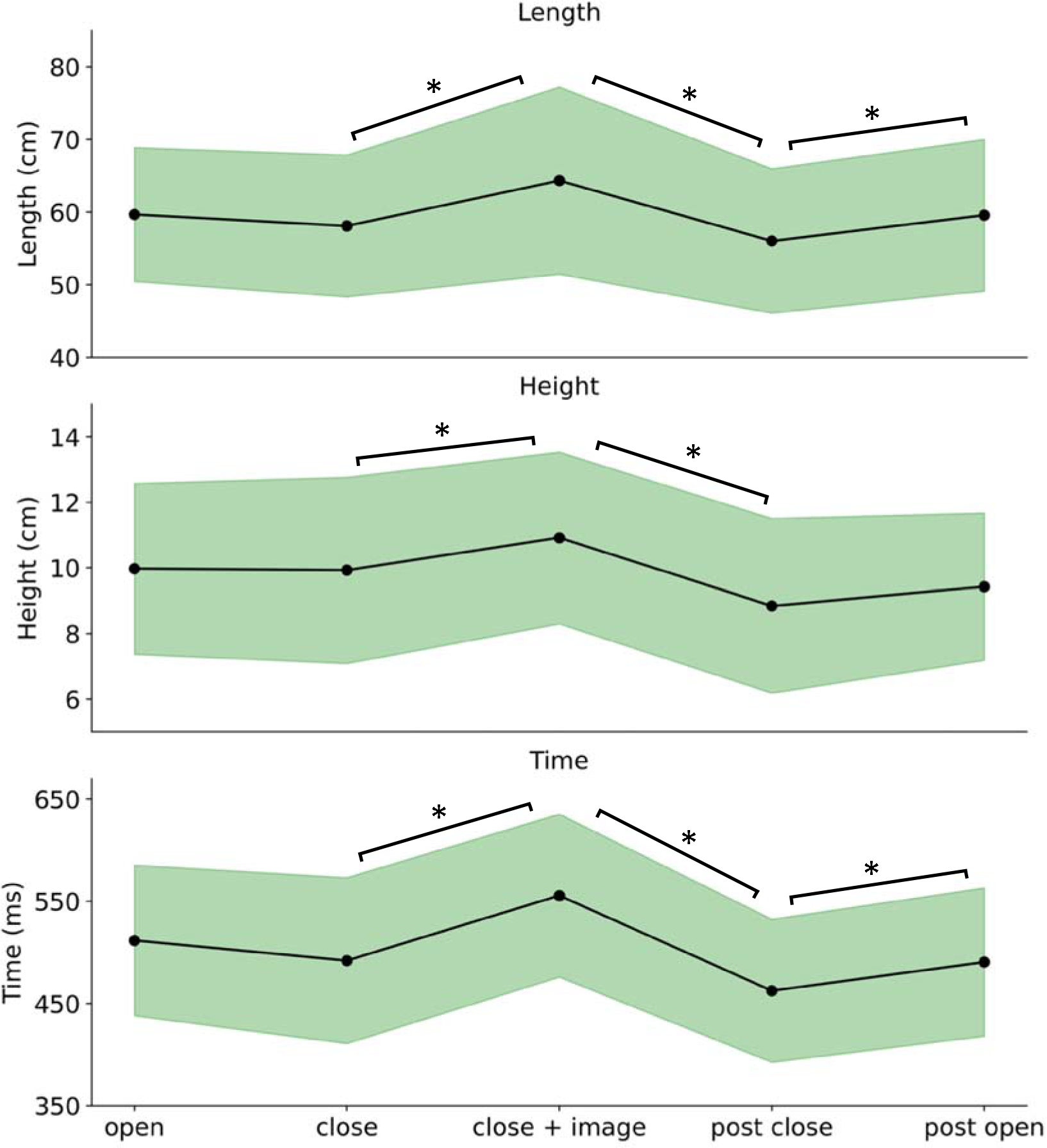
The average values of stride length, foot lift height, and time per step among participants for each condition. The bands represent the standard deviation. Significant differences were observed at p < 0.05.

The average height at which the participants lifted their feet for each condition is shown in Fig 2. A one-way analysis of variance (ANOVA) was conducted with conditions as the factors. This analysis revealed a significant main effect (*F* [4, 76] = 11.13, *p* = 3.70×10^-7^, η^*2*^ = 0.07). Subsequently, Bonferroni’s multiple comparisons test revealed significant differences between the closed eyes and closed eyes + image conditions as well as between the closed eyes + image and post-closed eyes conditions (p < 0.05).

The average time required for the first step for each condition is shown in Fig 2. A one-way analysis of variance was conducted with the conditions as the factors. This analysis revealed a significant main effect (*F* [4, 76] = 26.54, *p* = 8.67×10^-14^, η^*2*^ = 0.15). Subsequently, Bonferroni’s multiple comparisons revealed significant differences between the closed eyes and the closed eyes + image condition, between the closed eyes + image and the post-closed eyes condition, and between the closed eyes and the open eyes condition (p < 0.05).

## Discussion

The results of this study suggest that when participants assumed a large body image and starting to walk, the stride length, the time taken, and the height at which the foot is lifted increased. Previous studies have indicated that when the actual leg length is extended while maintaining the same body image, the stride length decreases [23]. In this study, the opposite phenomenon is considered to occur; to generate a torque larger than the actual body size, participants made a movement plan assuming a larger body image, resulting in an increases stride length. Increased vertical height was also observed to be necessary to advance the foot further.

Such changes are also believed to be influenced by the anticipated shift in the center of gravity. Our previous study has revealed the potential impact of the relative external environment on movement [34]. This impact may arise from the need to pre-adjust the body’s center of gravity when moving in various directions [35]. In this study, the participants’ movement output, assuming a larger shift in the center of gravity than in reality, is considered to be the result of a change in the movement plan.

In this study, a concept called “body schema” is used, which is similar to the “body image” concept. Body schemas can change owing to various factors. Body schema measurements sometimes make use of the peripersonal space (PPS). Peripersonal space refers to the existence of neurons that react when an object is near the body [36] and is believed to play a role in self-protection [37]. Previous studies have found that reaction speed increases when objects approach the body within the PPS, and the PPS has been defined as the space in which reaction speed increases [38]. PPS has been reported to expand body schemas by actively using tools and perceiving them as part of the body [39, 40]. Thus, although it has been reported that one’s own body schema is variable, the change is considered to occur unconsciously and is therefore believed to not affect the results of this study.

The results of this study do not support the findings of previous studies on walking movements during visual occlusion. Blocking visual information significantly affects walking movements. In a closed-eye state, the maximum vertical velocity of the center of mass was observed to be slower during the first step of walking [41]. Studies have also indicated that when perturbing the center of pressure (CoP) without providing feedback on CoP and visual target (field of view) during postural control, the disturbance of CoP increases [42]. Furthermore, regarding the role of visual information in postural control, no likely substitute for visual information has been suggested, as evidenced by increased disturbance of the CoP in the closed-eye state [43]. Additionally, when visual information is occluded during walking, studies have shown that variability in step width and duration increases regardless of walking speed, and particularly during slow walking, the instability becomes greater [24]. Other studies have reported that the duration of movement and step width increase to stabilize the center of mass [25]. However, the results of our study do not support the reported increase in step width under closed-eye conditions. One possible reason for this is that the walking movements in this study were performed barefoot. Walking barefoot has been reported to stabilize the body’s center of mass [44]. Therefore, the instability of the body’s center of mass reported in previous studies during closed-eye conditions did not occur in this study, resulting in no variation in the step width or movement between the open- and closed-eye conditions.

## Limitations and future prospects

A limitation of this study is that the instruction on body image was conducted through verbal instructions. Due to this verbal instruction, there may have been variations in the state of “body becoming larger” depending on the individual. Therefore, in the future, stable values can be obtained by investigating the motor output in a state where the body image is manipulated using self-aware stimuli, such as VR.

## Conclusion

In this study, we observed that when the body became larger, the stride length increased, resulting in an increase in time, and the height at which the foot was lifted also increased. Thus, our findings suggests that changing body image (cognition) could have an impact on actual movement.

